# Toxicogenomic identification of repositioned therapy for a monogenic disease

**DOI:** 10.1101/748863

**Authors:** Eric J. Kort, Nazish Sayed, Chun Liu, Sean M. Wu, Joseph C. Wu, Stefan Jovinge

**Author notes:** Correspondence to Stefan Jovinge.

## Abstract

The cost of drug development from initial concept to FDA approval has been estimated to be about 2.6 billion USD.^1^ This cost precludes development of targeted therapies for rare diseases such as monogenetic cardiomyopathies. As part of the Library of Integrated Network-based Cellular Signatures (LINCS) program funded by the NIH, the Broad Institute of MIT has publicly released transcriptional profiles quantifying the effects of more than 25,000 perturbagens on the expression of 978 genes in up to 77 cell lines.^2^ Transcriptomics has been shown to be a powerful tool in repurposing drugs^3,4^ and this dataset affords us the unique opportunity to systematically identify small molecule mimics or inhibitors of specific genes, thereby identifying novel treatments for genetic disorders. In this report, we take this approach to identify a novel drug therapy for a monogenic form of familial dilated cardiomyopathy with the transcriptional profile of FDA approved drugs. This approach could potentially be replicated for a wide range of monogenic diseases.

## Background

Mutations in the LMNA gene are associated with familial dilated cardiomyopathy^5-7^ as well as premature aging syndromes.^8^ LMNA encodes for Lamin A/C, a nuclear structural protein which also plays a key role in chromatin organization and transcriptional regulation.^9^ These mutations may be either missense mutations, leading to severe early disease presumably due to dominant negative or pathological gain of function, or nonsense mutations characterized by late-onset disease due to haploinsufficiency.^10,11^ LMNA has been shown to be involved in the regulation of several signaling pathways whose disruption may contribute to the disease phenotype. These include PDGF signaling, autophagy via Akt/Beclin signaling, MAPK signaling, and histone modification.^12-16^

## Results

In this study, we leveraged analysis of the LINCS L1000 dataset combined with iPSC based in vitro assays to identify FDA approved compounds that can reverse the effects of LMNA mutation at the transcriptional level (Fig. 1). We first obtained the Level 3 (normalized expression value) data from the NIH LINCS program. For our analysis, we utilized only the 978 directly measured genes (the “landmark” genes). The z-scores published by the LINCS program at the time we conducted this analysis were calculated for each sample vs. all the other samples on each experimental plate. For the purposes of our analysis, we wanted the z-scores calculated vs. controls only. Therefore, we calculated robust z-scores for each treated sample (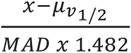 where 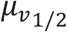 is the median value for the appropriate vehicle or empty vector controls on each plate).

**Figure 1.**
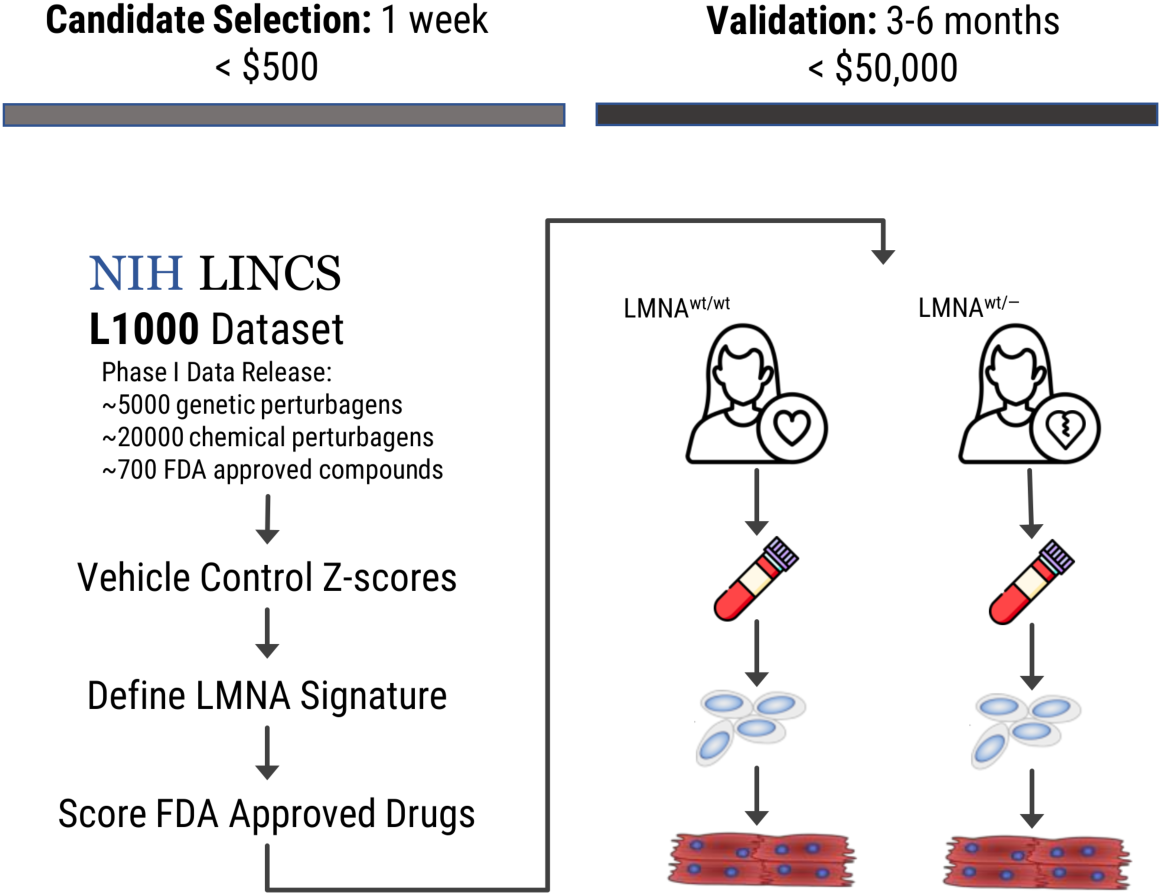
Overview of approach. The LINCS L1000 data was downloaded, and expression levels were converted to z-scores by comparing each treated sample to corresponding controls. From this dataset, we identify a gene-specific signature (in this case, the signature describing the transcriptional consequences of LMNA knockdown). With this signature, gene-targeting drugs are identified. The candidate drugs are then validated—in this case by testing in patient and healthy control iPSC-derived cardiomyocytes. Estimated duration and cost of this repositioning pipeline are provided for reference.

We next sought to identify those genes that *consistently* and *specifically* shifted in expression as a result of LMNA knockdown (Fig. 2). To find genes that consistently responded to LMNA knockdown, we performed a “rank of ranks” analysis. First, we ranked the z-scores (genes) within each sample that was treated with LMNA-targeting short hairpins (n=84) constructs in the LINCS dataset (Fig. 2A, left-hand panel). Next, we combined these ranks across the samples into a single list and tested each gene to see how random their ranks were across samples by means of the Kolmogorov Smirnov statistic—essentially inverting the traditional Gene Set Enrichment Analysis^17^ to quantify the enrichment of each gene across a set of samples rather than a set of genes within each sample (Fig. 2A, right hand panel).

**Figure 2.**
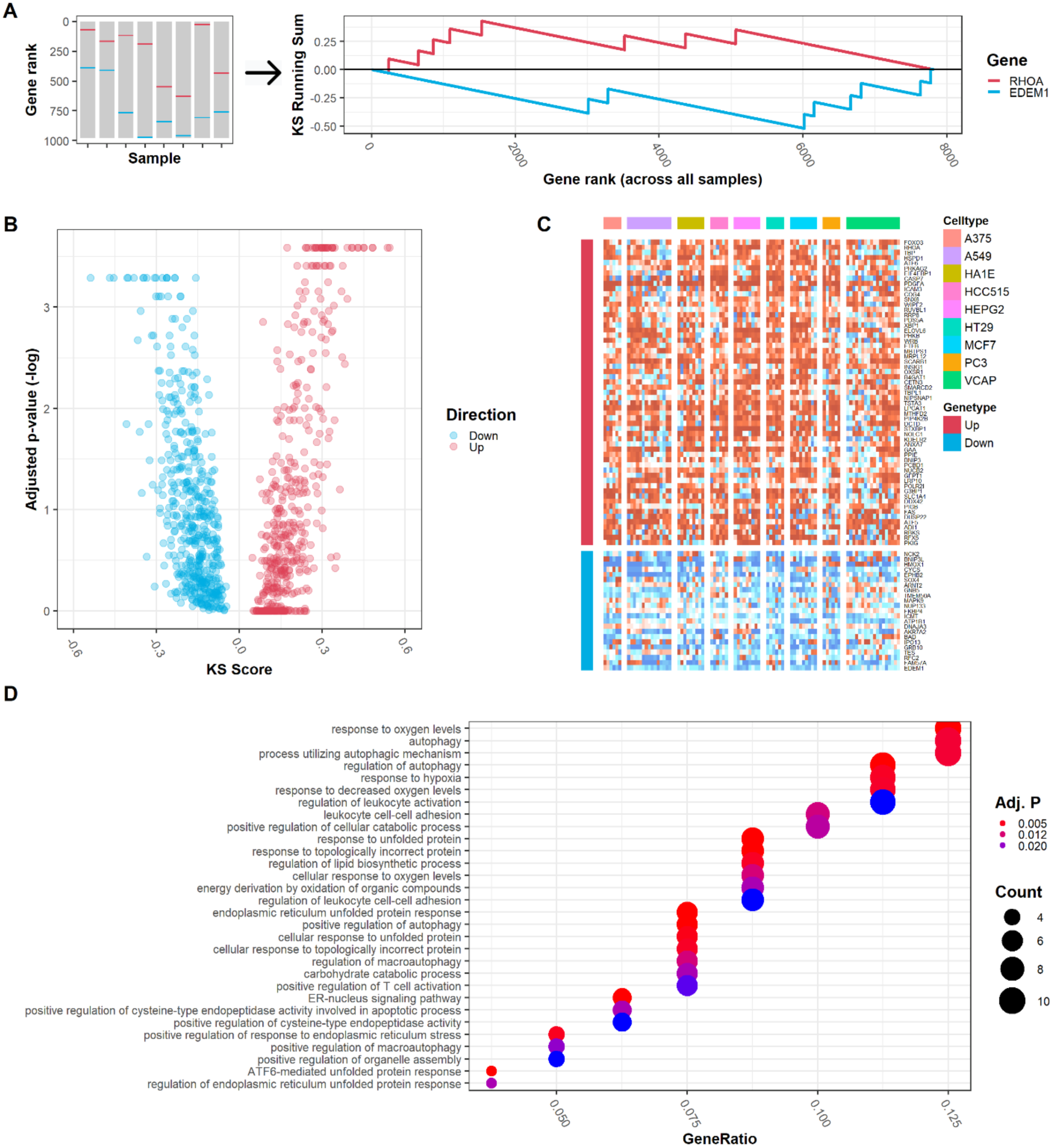
Establishment of LMNA knockdown gene signature. **(A)** Genes were scored based on how consistently and how strongly they were up- or down-regulated by LMNA knockdown in the LINCS L1000 database. Permutation of random sets of short hairpin perturbed samples was used to estimate the significance of these expression shifts. We selected those genes with both a KS score > 0.3 and a –log p-value > 0.3 as our LMNA knockdown gene set. **(B)** The LMNA knockdown gene signature showed fairly uniform up- or down-regulation across all samples in the L1000 dataset across all cell types treated. **(C)** Gene ontology enrichment analysis suggests that the genes in this signature are involved in pathways related to autophagy and apoptosis.

We also wanted to know how *specific* these gene expression changes were to LMNA knockdown vs. artefactual components of the experimental set up of the LINCS assay or non-specific cellular responses to genetic perturbation. Therefore, we repeated the above gene scoring using 100,000 random sets of gene-perturbed samples, with each set having the same number of samples as our LMNA perturbed set. This procedure allowed us to determine the probability that the differential expression statistic calculated for each gene from the LMNA perturbed samples was a unique feature of LMNA perturbation as opposed to a generic response frequently observed in random sets of gene-perturbed samples (Fig. 2b). The resulting bootstrapped p-values were adjusted to control the false discovery rate (FDR).^18^

Interestingly, while the most extreme differential scores were all highly significant as determined by this analysis, there were several genes that had intermediate differential expression scores that were nevertheless highly significant when compared to random perturbations (Fig. 2B). We selected those genes with FDR adjusted p-value < 0.001 and an enrichment (KS) score > 0.2 (absolute value) as our LMNA knockdown signature (supplemental Table S1). These genes were quite consistently down- or up-regulated across samples and cell lines (Fig. 2C).

PDGFA was among the most upregulated genes in our LMNA knockdown signature, consistent with recent work documenting activation of PDGF signaling in cardiomyocytes harboring mutation of LMNA.^12^ In addition, we performed Gene Ontology term enrichment analysis^19^ on the Biological Function Ontology for the genes in our signature (Fig 2C). This analysis identified multiple GO terms related to autophagy, apoptosis, and hypoxia response that were enriched in our LMNA knockdown gene set, also consistent with prior work.^7,20^

We next sought to identify drugs that could reverse this LMNA knockdown signature. There is some evidence that parametric approaches are quantifiably superior to non-parametric scoring metrics for this type of gene signature enrichment analysis. As a result, we chose the XSUM statistic to quantify reversal of the LMNA signature within drug treated samples.^21^ Since there were many samples per drug in the L1000 dataset, we took the median score for each drug. In this way, we informatically screened over 600 FDA approved drugs for their ability to reverse the LMNA knockdown signature. We again assigned p-values to these scores by permutation (based on 10,000 random gene signatures).

As validation, we submitted our LMNA signature to the original CMAP enrichment tool (https://portals.broadinstitute.org/cmap/)^22^ to see what drugs could reverse our knockdown signature as determined by the CMAP connectivity score (Table S2). Of the FDA approved drugs in our L1000 dataset, 6 demonstrated a significant ability to reverse the LMNA signature according the CMAP enrichment tool. Of these 6 drugs, 4 also had significant enrichment scores based on the XSUM statistic derived from the L1000 dataset (Fig. 3A). These 4 drugs had stronger enrichment scores based on the original CMAP tool compared to the 2 drugs that were not enriched in the L1000 analysis.

**Figure 3.**
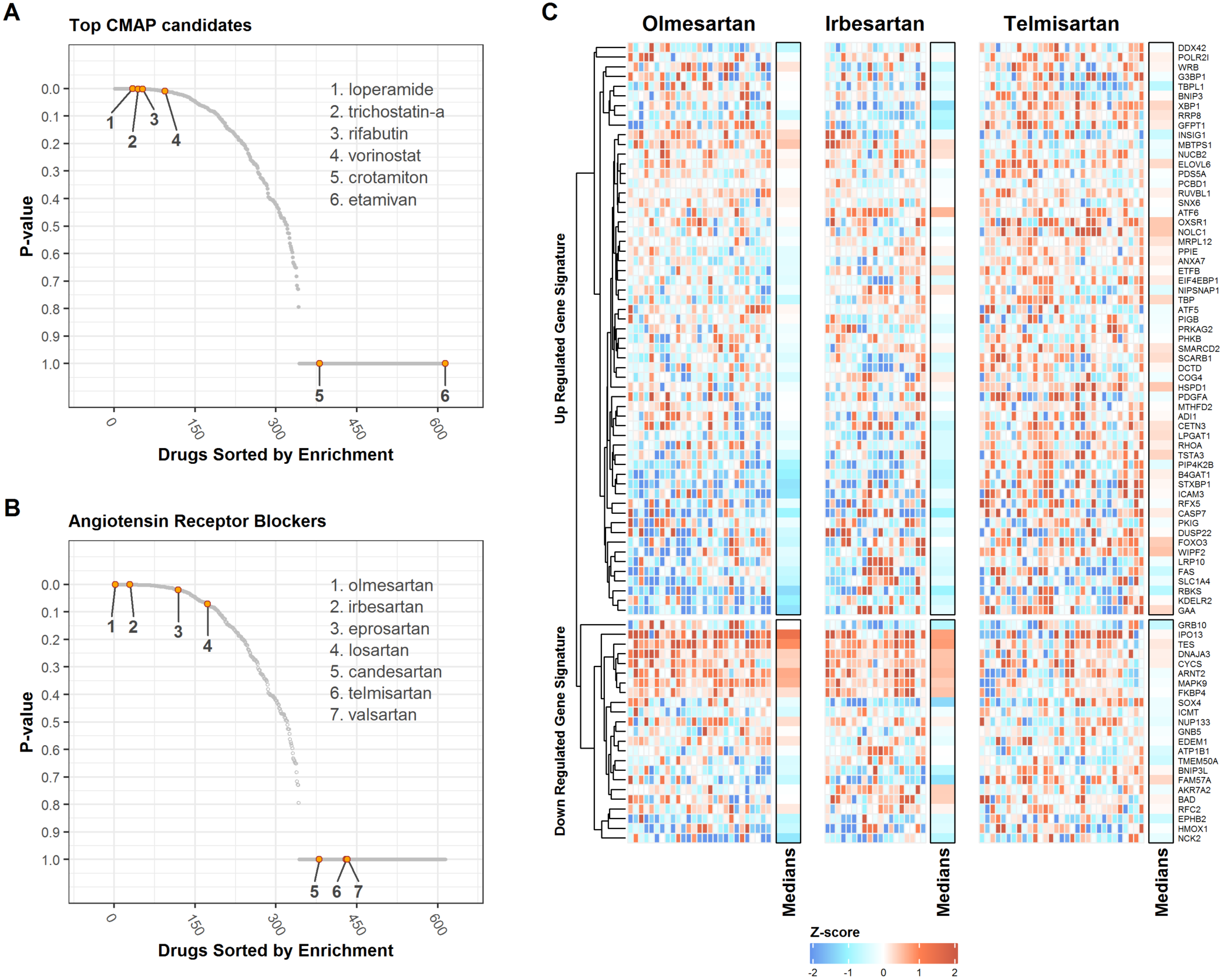
The LMNA knockdown signature was used to identify drugs that could reverse these transcriptional changes *in vitro*. **(A)** We first submitted the gene signature to the original CMAP web tool. Of the top 6 drugs that most strongly reversed the LMNA knockdown signature based on that tool, 4 also exhibited highly significant reversal of this signature in the LINCS L1000 data, suggesting there was some coherence between the various platforms and analytic approaches used by these systems. **(B)** We note that two angiotensin receptor blockers were among the most highly significant drugs based on their ability to reverse the LMNA knockdown signature. Indeed, 3 of 7 ARBs in the database showed gene set enrichment with an FDR adjusted p-value of < 0.05. **(C)** These ARBs do not reverse the entirety of the LMNA signature. Rather, high scoring ARBs (olmesartan and irbesartan) reverse highly overlapping segments of the LMNA signature, whereas telmisartan (a low scoring drug) exhibits a more random expression pattern for the genes in the LMNA knockdown signature.

We noted that multiple angiotensin receptor blockers (ARBs) exhibited significant reversal of the LMNA knockdown signature (Table S3 and Fig 4B), including olmesartan which was among the group of drugs that reversed this signature most significantly. However, not all ARBs exhibited this feature. It has previously been demonstrated that the various members of this drug class have variable transcriptional effects.^23-25^ When we analyzed all ARBs in our dataset as a group, this class of drugs collectively exhibited a significant reversal of the LMNA knockdown signature, but this effect was weaker for the entire class than for olmesartan or irbesartan specifically (FDR = 0.0147 vs. 0.0001 and 0.0021, respectively).

**Figure 4.**
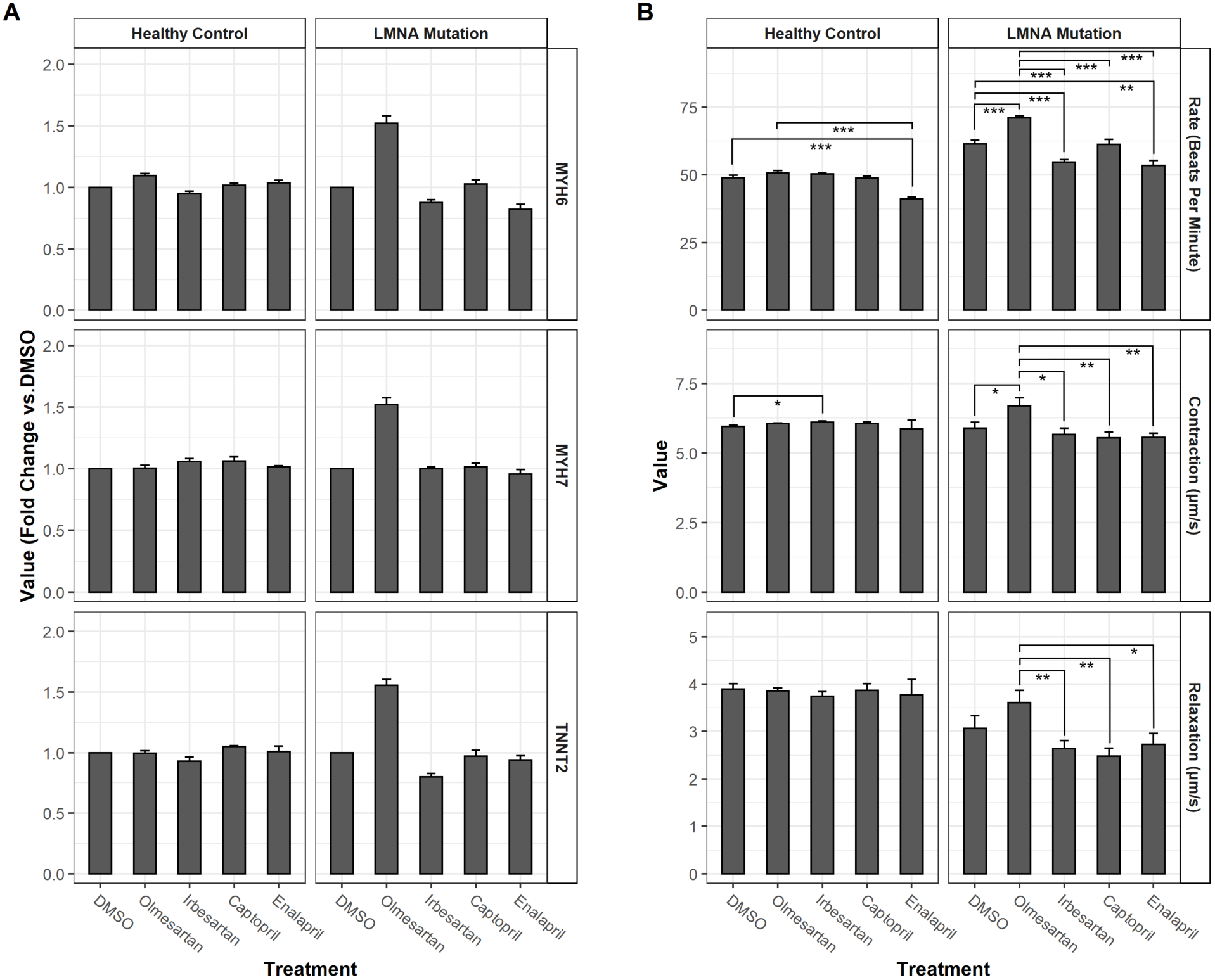
In vitro validation of the effect of predicted drugs on cardiomyocyte transcription and phenotype. All results are from duplicate experiments, with 3 technical replicates per experiment (total N=6). All error bars are +SEM. **(A)** We measured the expression of three cardiac markers (MYH6, MYH7, and TNNT2) by quantitative PCR in cardiomyocytes derived from iPSC cells from a healthy control and a patient harboring a LMNA mutation. Fold-change relative to DMSO treated controls was calculated for each marker after 48 hours of treatment with the drugs shown, which include 2 of the ARBs identified by our analysis, and two angiotensin converting enzyme inhibitors used as controls due to their canonical activity on the same pathway as ARBs. Significant differences from DMSO treated samples are indicated (*: p ≤ 0.05, **: p≤ 0.01, ***: p≤0.001 by two-tailed t-tests). **(B)** Beating rate, contractile velocity, and relaxation velocity were measured by quantitative video microscopy in both control and LMNA mutant cardiomyocytes. Rate is expressed in beats per minute. Contractile force is expressed in micrometers per second. Significant differences from DMSO and/or olmesartan treated samples are indicated (*: p ≤ 0.05, **: p≤ 0.01, ***: p≤0.001 by two-tailed t-tests).

When we examined the effect of the highest scoring ARBs (olmesartan and irbesartan) on the expression of the individual genes in the LMNA knockdown signature, we noted that these drugs down regulated many of the genes upregulated by LMNA knockdown, and vice versa (Fig 4C). There was no such trend apparent for telmisartan, which did not exhibit significant enrichment in our analysis. However, the transcriptional effects of olmesartan and irbesartan were not universal with respect to the LMNA regulated genes, indicating that these drugs were acting on only portions of the LMNA knockdown “axis”.

We next tested whether olmesartan and/or irbesartan treatment could influence the expression of cardiac markers and/or function of cardiomyocytes derived from induced pluripotent stem cells (iPSC-CMs) harboring a disease-related LMNA nonsense mutation. These cells harbor a heterozygous insertion of a guanine between nucleotides 348 and 349, causing a frameshift mutation at codon 117, and resulting premature stop at codon 129. In iPSC-CMs from a healthy control patient treated for 48 hours, a slight increase in MYH6 was observed in olmesartan treated cells, and a slight increase in TNNT2 expression was observed in captopril treated cells (Fig. 4A). In contrast, there was a 1.5-fold increase in MYH6, MYH7, and TNNT2 in olmesartan treated iPSC-CMs derived from a patient harboring a nonsense LMNA mutation. Other treatments tested either had no effect on these markers or suppressed their expression. We hypothesized that increased expression of these sarcomeric genes would correspond to improved contractility in these LMNA mutant cells.

To test that hypothesis, we examined what effect, if any, treatment of iPSC-CMs with these drugs had on the functional properties of these cells. For this, we examined the contractile properties of healthy control and LMNA-mutated iPSC-CMs. Irbesartan treatment of iPSC-CMs derived from a control patient produced a very small (though statistically significant) increase in contraction velocity (Fig 4B), but no significant effect on contractile rate relative to DMSO treated controls, while olmesartan had no effect on the behavior of these cells relative to DMSO control. In contrast, olmesartan treatment of iPSC-CMs derived from the LMNA-mutation patient resulted in a marked increase in both contraction velocity and beating rate. While there was a trend towards increased relaxation velocity in olmesartan treated cells compared to DMSO only control, this difference was not significant. However, olmesartan treatment was associated with significantly faster relaxation velocity as compared to the other drugs tested. These observations support the hypothesis that olmesartan will not exacerbate any underlying diastolic dysfunction and may in fact be favorable to other heart failure medications with respect to diastolic function.

## Conclusions

These results demonstrate that the LINCS L1000 database can be exploited to perform an *in silico* screen for repositionable drugs that reverse the transcriptional consequences of specific genetic perturbations. One of two treatments we identified in such a way for LMNA mutation related cardiomyopathy showed a favorable response in our *in vitro* model—suggesting that this approach to drug repositioning for rare diseases may be a promising alternative to the traditional drug development pipeline.

## Methods

### Statistics

No statistical methods were used to predetermine sample size. The experiments were not randomized. The investigators who performed the phenotype assessment of the CMs were blinded to group allocation during experiments and data collection. The studies comply with all ethical regulations.

### Data Availability

This analysis was performed on the initial release for the LINCS L1000 dataset. Updated data from LINCS is available under accession #GSE92742. The relevant same plate vehicle/vector control z-scores we calculated from the original LINCS L1000 data release, as well as R scripts to reproduce the analysis and figures presented here, are available at https://github.com/vanandelinstitute/Lamin.

### LMNA Gene Signature Definition

Full details of the generation of the LMNA gene expression signature and drug selection based on that signature using the LINCS L1000 database, are provided in a supplementary file (data_analysis.html). This information and supporting data required to repeat the analysis presented in this paper (including regenerating the figures) are freely available from github: https://github.com/vanandelinstitute/Lamin. Briefly, normalized gene expression data was obtained from the LINCS L1000 program. We then calculated the robust z-score for each gene within each sample relative to vehicle treated samples of the same cell type on the same 384 well plate. We extracted the z-scores for all instances treated with short hairpins targeting LMNA. This data was ranked sample-wise. Second, the entire matrix of ranks (978 genes by 84 shRNA samples) was ranked and Kolmogorov Smirnov analysis was performed on the position of each occurrence of each gene within this vector of ranks. The resulting analysis quantifies the extent to which the expression of each gene was consistently biased up or down relative to all other genes.

Finally, a bootstrapping procedure as performed to estimate significance of the KS score for each gene. We calculated KS scores for 100,000 random sets of shRNA treated samples (84 samples in each set to match the LMNA set) in the LINCS data. The resulting p-values were adjusted for multiple comparisons using the method of Benjamini and Hochberg to control the false discovery rate at less than 5%.^18^

### Drug Selection

To estimate the bias in gene expression for the genes in our LMNA signature within each sample in the L1000 dataset treated with and FDA approved compound, we used the XSUM metric because there is some evidence it is among the more performant algorithms for CMAP type data.^21^ The XSUM limits its search to the top N variable genes. However, since the L1000 dataset is already confined to the 978 most variant genes in the genome as determined by the LINCS program, we did not filter the gene set further. Therefore, we take the sum of the z-scores for our upregulated genes and subtract the sum of the z-scores of the down regulated genes within the 978 L1000 genes for each drug perturbed instance. Because there are multiple instances per drug, we collapsed these scores to a single score per drug by taking the median.We again used a bootstrapping procedure to estimate the significance of each drug’s score relative to random perturbations. We scored 10,000 random gene signatures (each with the same number of “up” and “down” regulated genes as the LMNA signature) to estimate how specific each drug was to the LMNA signature. Drugs were then ranked based on their bootstrapped p-value.

### Generation of human iPSCs

Protocol for isolation and use of patient blood-derived peripheral blood mononuclear cell (PBMC) were approved by the Stanford University Human Subjects Research Institutional Review Board. PBMCs were isolated using a Ficoll-Paque PLUS gradient (GE Healthcare) and expanded as previously reported.^26^ For reprogramming, 1 million PBMCs were plated in medium supplemented with four OSKM reprogramming factors (CytoTune-iPS Sendai Reprogramming Kit, Life Technologies) according to the manufacturer’s recommendations. The medium was changed after 24hr transfection and transferred to E7N medium (E8 medium minus TGFβ1 and 200 µM sodium butyrate) on day 3. Colonies were picked into 1 well of a 12-well plate (1-colony in each well) on around day 20 and cultured in E8 with 10 µM Y-27632 (Selleckchem). hiPSCs were then expanded into 6-well plated (coated with 1:200 growth factor-reduced Matrigel) and maintained in E8 medium. Confluent hiPSCs were passaged every four days using 0.5 mM EDTA.

### Differentiation of hiPSCs to cardiomyocytes

hiPSCs were routinely maintained in 6-well plates as described above. Cells were grown to reach 90% confluency and then subjected to differentiation in RPMI/B27 without insulin medium (Life Technologies) supplemented with 6 µM CHIR99021 (Selleckchem). Following 48h, the cells were subjected to the same medium supplemented with 4 µM IWR-1-endo (Selleckchem). On day 7, the medium was changed to RPMI-B27 with insulin and exchanged every other day. Beating hiPSC-CMs usually can be observed around day 7 to day 10. On day 11, the medium was switched to RPMI-B27 without D-glucose (Life Technologies) for 4 days to purify cardiomyocytes. For drug treatment and function analysis, purified iPSC-CMs were dissociated using TrypLE Express (Life Technologies) and re-plated to Matrigel-coated plates accordingly.

### Drug Treatment

The indicated drug compounds were reconstituted from powder in DMSO to a working concentration of 10mM. Drugs were then added to the cell culture wells to a final concentration of 10μM and the cell were returned to the cell culture incubator for the indicated times. Controls were treated with DMSO alone.

### Quantitative real-time PCR

RNA was extracted using a QIAGEN RNeasy kit following the manufacturer’s instructions. cDNA was synthesized from 100 ng of total RNA using the High Capacity RNA-to-cDNA kit (ThermoFisher Scientific). Realtime-PCR was performed using TaqMan Gene Expression Master Mix and TaqMan probes (GAPDH, Hs02758991_g1; TNNT2, Hs00165960_m1; MYH6, Hs01101425_m1; MYH7, Hs01110632_m1). PCR reactions were conducted on 7900HT Real-Time PCR system (ThermoFisher Scientific) with triplicates and assessed using ΔΔCt relative quantification (RQ) method normalizing to GAPDH housekeeping gene.

### High-content video-based cardiomyocyte contractility analysis

hiPSC differentiated cardiomyocytes were plated onto Matrigel-coated 96 well plates (40,000 per well) as described above. Following treatment with drugs, iPSC-CMs were examined on Sony SI8000 Live Cell Imaging System (Sony Biotechnology) with CO_2_ and 37 °C temperature incubation. Cell activities were recorded the beating video at a high frame rate (150 fps), focus and light conditions were automated controlled by the SI8000 software. After data acquisition, displacement and magnitudes of cardiomyocyte motions were calculated and presented using a motion detection algorithm by SI8000 software.

## Acknowledgement

This work was mainly funded through Richard and Helen DeVos Foundation

---

**Supplemental Table S1:**
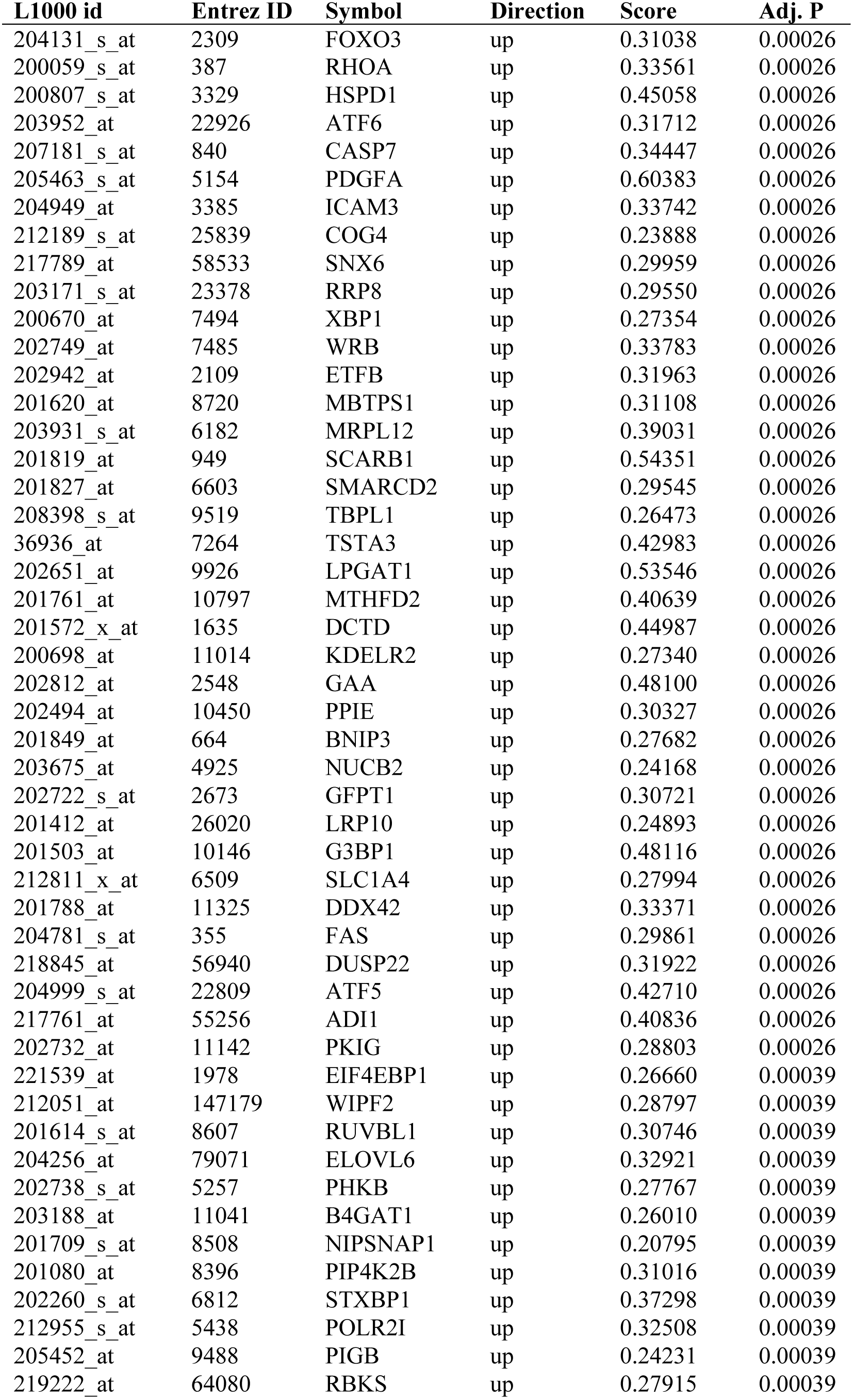
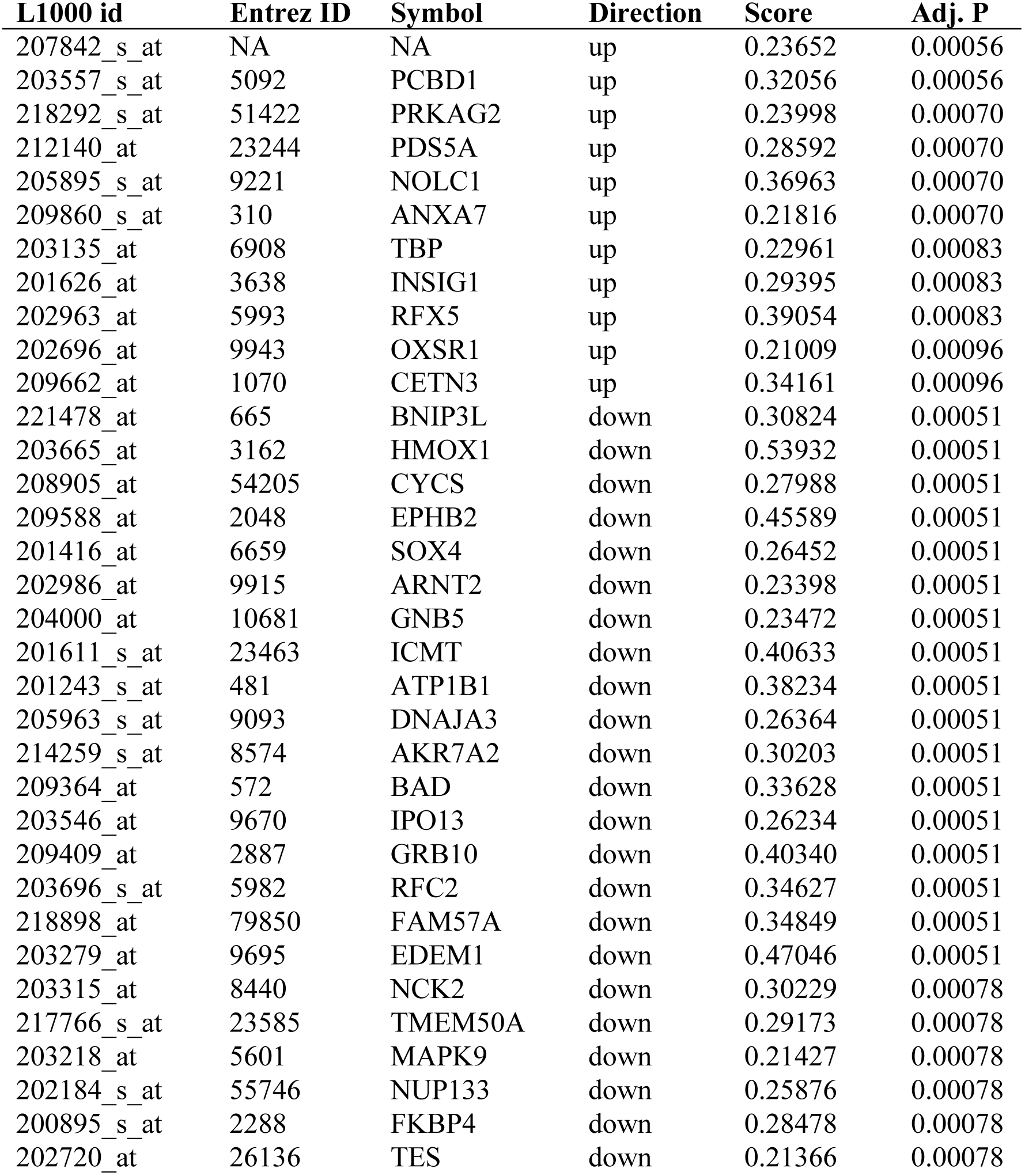
LMNA Knockdown Signature Genes. Genes identified as consistently and specifically perturbed by LMNA targeting short hairpin treatment. The direction column indicates whether the gene is up-regulated or down-regulated by short hairpin treatment relative to controls.

**Supplemental Table S2:** Drugs which significantly reverse the LMNA knockdown signature as determined by the original CMAP web tool.

| rank | cmap name | mean | n | enrichment | p |
| --- | --- | --- | --- | --- | --- |
| 1 | vorinostat | -0.599 | 12 | -0.771 | 0 |
| 2 | trichostatin A | -0.412 | 182 | -0.5 | 0 |
| 3 | scriptaid | -0.684 | 3 | -0.941 | 0.00032 |
| 4 | rifabutin | -0.608 | 3 | -0.889 | 0.00268 |
| 5 | rolitetracycline | -0.247 | 4 | -0.801 | 0.00304 |
| 7 | difenidol | -0.381 | 3 | -0.859 | 0.00563 |
| 9 | methacholine<br>chloride | -0.407 | 3 | -0.851 | 0.00659 |
| 13 | levamisole | -0.325 | 4 | -0.728 | 0.01112 |
| 15 | loperamide | -0.438 | 6 | -0.603 | 0.01357 |
| 16 | sulfadiazine | -0.322 | 5 | -0.645 | 0.01402 |
| 18 | lobelanidine | -0.35 | 4 | -0.706 | 0.01554 |
| 19 | GW-8510 | -0.445 | 4 | -0.701 | 0.01649 |
| 20 | calmidazolium | -0.542 | 2 | -0.906 | 0.01795 |
| 21 | ticarcillin | -0.487 | 3 | -0.775 | 0.02342 |
| 23 | MK-886 | -0.483 | 2 | -0.882 | 0.02813 |
| 24 | imipenem | -0.358 | 4 | -0.665 | 0.02847 |
| 27 | suloctidil | -0.339 | 4 | -0.651 | 0.03495 |
| 29 | crotamiton | -0.43 | 4 | -0.644 | 0.03847 |
| 31 | piracetam | -0.335 | 4 | -0.636 | 0.04269 |
| 33 | etamivan | -0.313 | 4 | -0.633 | 0.04434 |

**Supplemental Table S3:**
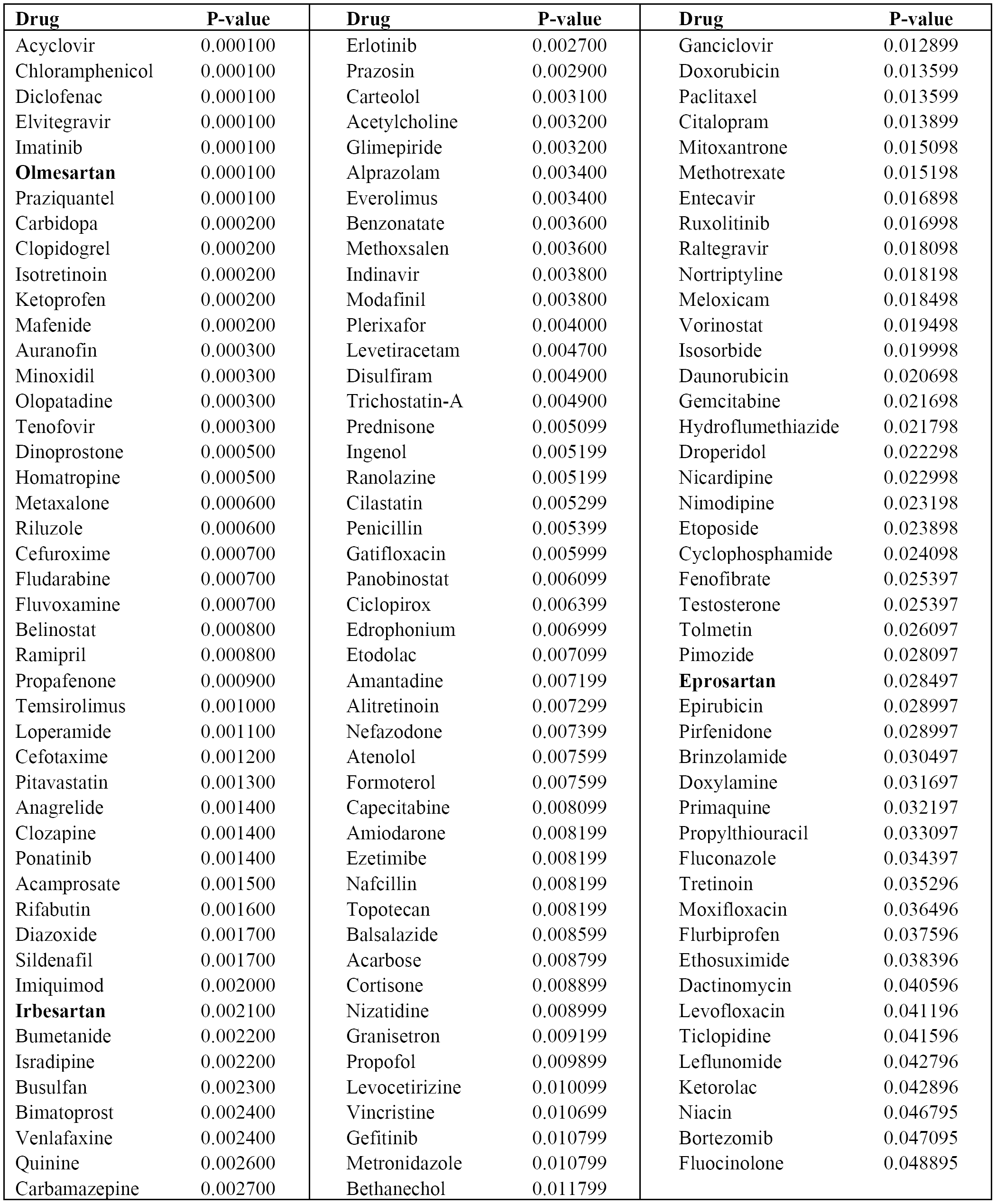
Drugs which significantly reversed LMNA knockdown signature in our analysis of L1000 dataset.

